# Whole genome analysis of the methylome and hydroxymethylome in normal and malignant lung and liver

**DOI:** 10.1101/062588

**Authors:** Xin Li, Yun Liu, Tal Salz, Kasper D. Hansen, Andrew Feinberg

## Abstract

DNA methylation at the 5-postion of cytosine (5mC) is a well-established epigenetic modification which regulates gene expression and cellular plasticity in development and disease. The ten-eleven translocation (TET) gene family is able to oxidize 5mC to 5-hydroxymethyl-cytosine (5hmC), providing an active mechanism for DNA demethylation, and may also provide its own regulatory function. Here we applied oxidative bisulfite sequencing to generate whole-genome DNA methylation and hydroxymethylation maps at single-base resolution in paired human liver and lung normal and cancer. We found that 5hmC is significantly enriched in CpG island (CGI) shores while depleted in CGIs themselves, in particular at active genes, resulting in a 5hmC but not 5mC bimodal distribution around CGI corresponding to H3K4me1 marks. Hydroxymethylation on promoters, gene bodies, and transcription termination regions showed strong positive correlation with gene expression within and across tissues, suggesting that 5hmC is a mark of active genes and could play a role gene expression mediated by DNA demethylation. Comparative analysis of methylomes and hydroxymethylomes revealed that 5hmC is significantly enriched in both tissue specific DMRs (t-DMRs) and cancer specific DMRs (c-DMRs), and 5hmC is negatively correlated with methylation changes, particularly in non-CGI associated DMRs. Together these findings indicate that changes in 5mC as well as in 5hmC and coupled to H3K4me1 correspond to differential gene expression in tissues and matching tumors, revealing an intricate gene expression regulation through interplay of methylome, hydroxyl-methylome, and histone modifications.

## Introduction

DNA methylation is an important epigenetic modification which plays a role in diverse biological processes, including maintenance of genomic stability, gene silencing, embryonic development, and tumorigenesis (Jones 2012; Smith and Meissner 2013; Timp and Feinberg 2013). Recently, a family of Ten-eleven translocation methylcytosine dioxygenase (TET) proteins was shown to oxidize 5mC (5-methylcytosine) to 5hmC (5-hydroxymethylcytosine) (Laird et al. 2013; Wu and Zhang 2014). TET-mediated 5hmC is abundant in variety of mammalian tissues, such as brain and stem cells (Pastor et al. 2013), and plays a role in in epigenetic reprogramming, cell differentiation and tumorigenesis.

In contrast to a relatively constant 5mC level across tissues, 5hmC is highly tissue-specific (Nestor et al. 2012). 5hmC has been shown to be enriched in promoters, gene bodies, and distal cis-regulatory elements, such as enhancers, and thus potentially involved in regulation of tissue-specific gene expression (Pastor et al. 2013; Wu and Zhang 2014). However, how 5hmC regulates and shapes tissue-specific epigenomes through DNA demethylation remains largely unknown.

Significant global loss of hydroxymethylation has been observed in cancer, and disruption of TET-mediated DNA demethylation was proposed to contribute to tumorigenesis (Kudo et al. 2012; Pfeifer et al. 2013). Loss-of-function mutations in TET2 are found in myelodysplastic syndrome (Delhommeau et al. 2009; Langemeijer et al. 2009), and mutations in the isocitrate dehydrogenase IDH1/IDH2 genes, co-factors of TET enzymes, are found in glioma, acute myeloid leukemia, and melanoma (Dang et al. 2010; Shibata et al. 2011). Down regulation of TETs and IDHs is also found in several cancer types (Lian et al. 2012; Yang et al. 2013). By depletion of TET enzymes in a pluripotent embryonic carcinoma cell (ECC) model, it was proposed that disruption of TET-mediated 5hmC, which associated with transcriptionally active chromatin environment, represented a crucial step toward aberrant gene silencing through DNA methylation in cancer cells (Putiri et al. 2014). Limited by the lack of high-resolution hydroxymethylomes in normal and matched-tumor tissues, dynamic changes in 5hmC and underlying mechanisms during carcinogenesis have not yet been elucidated.

Distinguishing 5hmC from 5mC is virtually impossible by traditional bisulfite conversion based methods. Several strategies have been developed to label and enrich 5hmC followed by sequencing with limited resolution (Laird et al. 2013; Putiri et al. 2014). Recently two methods were implemented allowing for quantification of 5hmC at a single-base resolution. TET-assisted bisulfite sequencing (TAB-Seq) (Yu et al. 2012) makes use of enzymatic reactions; β-glucosyltransferase to glucosylate 5hmC and TET1 for subsequent oxidation to produce 5-carboxylcytosine (5caC). In the final step of bisulfite treatment both 5caC and unmodified cytosine are converted to uracil allowing the identification of 5hmC, which is read as cytosine during sequencing. Oxidative bisulfite sequencing (oxBS-Seq) (Booth et al. 2012) rather makes use of the highly selective chemical oxidant potassium perruthenate (KRuO_4_) to convert 5hmC to 5-formylcytosine (5fC), followed by bisulfite conversion and sequencing. This library is then compared to a traditional bisulfite sequencing library (BS-Seq) constructed on the same sample to identify differences in 5mC which accounts for 5hmC positions. oxBS-Seq has a relatively simple and fast experimental workflow and can obtain both methylome and hydroxymethylome simultaneously.

Here, we utilized the oxBS-Seq method to study, at single base-resolution, the methylomes and hydroxymethylomes of a normal and matching tumor set from human lung and liver tissues, providing a valuable resource for studying 5hmC landscapes in different tissues and understanding the roles of 5hmC during tumorigenesis. Integrated with RNA-Seq and ChIP-Seq data, we described the genome-wide hydroxymethylation pattern in normal and tumor tissues, and then investigated the relationship among 5mC, 5hmC and histone modifications. Finally, analysis of differentially methylated regions (DMRs) in normal and tumor tissues revealed a negative correlation between changes of 5mC and 5hmC/H3K4me1. Together our findings demonstrated an intricate gene expression regulation through interplay of methylome, hydroxyl-methylome, and histone modifications during tissue differentiation and tumorigenesis.

## Results

### Generation of single-base resolution hydroxymethylation and methylation maps in matched human normal and tumor samples from lung and liver

We applied oxBS-Seq and BS-Seq to genomic DNA extracted from four human liver normal-tumor pairs and three human lung normal-tumor pairs (14 samples total). All libraries were sequenced to an average depth of 15.4 × per CpG cytosine (Supplemental Table S1). In order to evaluate bisulfite (BS) and oxidative bisulfite (oxBS) conversion rates, non-methylated *E. coli* and CpG hydroxymethylated lambda phage DNA were spiked in as controls during library preparation. Based on spike-in controls, both high bisulfite (unmethylated cytosine to uracil, 99.66%) and high oxidative bisulfite conversion (5-hydroxymethycytosine to uracil, 96.57%) were observed (Supplemental Table S1).

According to the oxBS-Seq method principle (Booth et al. 2013), hydroxymethylation level of cytosines was calculated for each sample based on the differential methylation between oxBS-Seq and the corresponding BS-Seq libraries. Since human tissues, except for brain, exhibit relatively low abundance of 5hmC (Nestor et al. 2012), high sequencing coverage was suggested in order to achieve an accurate 5hmC measurement at single CpG sites (Booth et al. 2013). Here, in order to confidently identify 5hmC-enrich regions and sites, we took advantage of the information from biological replicates and adjacent CpG sites, using an algorithm based on local likelihood smoothing (BSmooth) to identify consensus 5hmC regions and sites for each normal and tumor group. This method was successfully applied to DMR identification between groups with low sequencing coverage in our previous studies (Hansen et al. 2011; Hansen et al. 2014).

In total 89,437 to 297,724 5hmC regions containing 1,013,839 to 3,178,223 CpG sites were identified in normal tissues, while only 2,255 to 37,159 5hmC regions containing 15,567 to 365,443 CpG sites in matching tumor tissues (Supplemental Table S2-6). The global hydroxymethylation levels of normal tissues are 2.27 to 5.68% in liver and 1.94 to 3.04% in lung, while significantly lower in the matched tumors (liver: 0.70 to 2.07%; lung: 0.65 to 1.07%). These results are similar to those achieved by mass-spectrometry or antibody-based assays (Jin et al. 2011; Lian et al. 2012; Yang et al. 2013). Considering incomplete conversion of 5hmC to 5fC in oxBS-seq that could result in false negative of 5hmC detection, the reported global 5hmC level here is a conservative underestimation. In contrast to the small variation of global 5mC level among patients in normal tissues, global 5hmC level showed considerable variation in both normal and tumor tissues (Supplemental Fig. S1).

To further verify our BSmooth method, we did deep sequencing (55 × coverage) for one liver normal sample and compared the estimated 5mC and 5hmC profiles based on 55 × data to that based on original 15 × data. High correlations (5mC: 0.96 and 5hmC: 0.82) were observed between two sets to data when smoothing both high and low coverage data (Supplemental Fig. S2). Additionally, we estimated 5mC and 5hmC levels based on high coverage data over 2 kb interval using only CpGs at least 20 × coverage across whole genome and compared them to smoothed results based on low coverage data. We also found close agreement between them for both 5mC (0.90) and 5hmC (0.79). These results validated our BSmooth approach that estimate 5mC and 5hmC levels with high accuracy using relatively low coverage data. Furthermore, 15 identified liver c-DMRs and 5hmC regions were picked out, and all 15 c-DMRs and 13 out of 15 5hmC regions were replicated using another cohort including 6 liver normal-cancer pairs (Supplemental Fig. S3-4), again confirming high accuracy of our approach for identifying both DMRs and 5hmCs.

### Hydroxymethylation landscapes of human normal and malignant liver and lung

To explore the hydroxymethylation landscape, we first examined genomic distribution of 5hmC in each tissue according to annotated genomic features. Compared to the whole genome 5hmC average level, 5hmC was depleted in regions around transcriptional start sites (TSSs) and in intergenic regions. In contrast, 5hmC was highly enriched at enhancers, especially active enhancers, in all tissues (Fig. 1A), which is consistent with previous studies in stem cells and brain tissue (Yu et al. 2012; Wen et al. 2014). Notably, 5hmC was highly enriched at CGI shores while depleted at CGIs, which are largely overlapped with TSSs (Fig. 1A). Our and others’ previous studies showed that both cancer DMRs (c-DMRs) and tissue DMRs (t-DMRs) are mostly located at CGI shores rather than CGIs, and that differential DNA methylation at CGI shores strongly correlates with differential gene expression when comparing different tissues as well as normal and tumor tissues (Irizarry et al. 2009; Hansen et al. 2011; Pathiraja et al. 2014). Together, those findings suggest that 5hmC could play a particular part in regulating DNA methylation at CGI shores.

**Figure 1.**
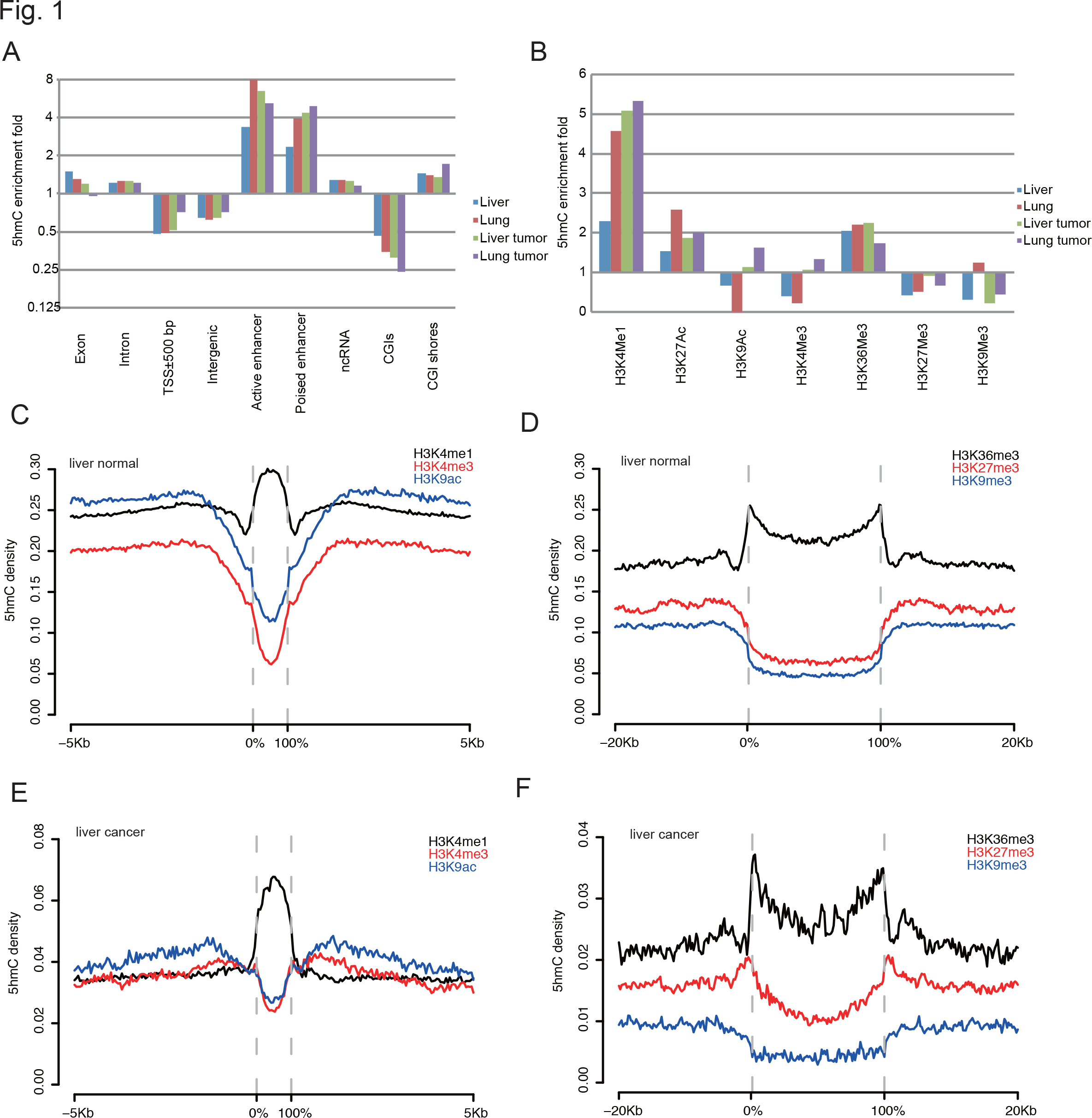
The hydroxymethylation landscapes of human normal tumor tissues. (A-B) 5hmC fold enrichment in different genomic features (A) and histone modifications (B). 5hmC enrichment fold in each genomic feature was calculated as the ratio of 5hmC density of that feature to genome-wide average 5hmC density. H3K9Ac ChIP-Seq data for lung normal is not available. (C-F) 5hmC density distribution across narrow and broad histone modifications and their flanking regions in liver normal (C, E) and liver tumor samples (D, F). 0% and 100% x-axis indicate start and end of all called histone peaks/domains. **For each liver/lung normal/cancer group, 5hmC level/density among replicates was averaged among replicates.**

Next, we examined the relationship between 5hmC and chromatin organization, using chromatin immunoprecipitation sequencing (ChIP-Seq) data from the Roadmap Epigenomics Project (Roadmap Epigenomics et al. 2015). In general, 5hmC positively correlated with active chromatin features, including active histone modification (H3K4me1/2, H3K27ac and H3K36me3) and DNase hypersensitive sites, with the exception of promoter-associated marks (H3K4me3 and H2A.Z) (Fig. 1B-F and Supplemental Fig. S5). On the other hand, 5hmC showed negative correlation with repressive chromatin marks, such as H3K27me3 and H3K9me3 (Fig. 1B-F and Supplemental Fig. S5). Our analysis of 5hmC and chromatin organization further support other studies indicating that 5hmC is an active epigenetic mark that corresponds to open chromatin (Wen et al. 2014).

Large hypomethylated blocks and small DMRs associated with loss of CGI methylation boundary stability is a common feature of the cancer epigenome (Timp and Feinberg 2013). To explore the role of 5hmC in DNA methylation change, we first examined the distribution of 5hmC among different cancer DMR categories. It is noteworthy that previous studies using traditional bisulfite sequencing method cannot distinguish 5mC and 5hmC. Therefore, the previously identified DMRs are mixture of both methylation and hydroxymethylation differences. Using data from oxBS-Seq, we were able to identify DMRs by only taking methylation changes into account between normal and matched tumor samples. Identified DMRs were grouped into small-scale DMRs and large-scale blocks according to their length (see Methods). We found that in both liver and lung c-DMRs, 5hmC is enriched in hypermethylated blocks and small DMRs, but not in hypomethylated blocks (Supplemental Fig. S6A-B). We also compared the distribution of 5hmC among different t-DMR groups between normal liver and lung tissues and observed similar pattern (Supplemental Fig. S6C). This result suggests that 5hmC and the responsible DNA demethylation pathway might be involved for methylation regulation on small-scale DMRs, while large-scale genomic hypomethylation may have different underlying mechanisms not involving active DNA demethylation.

### 5hmC distribution around genic regions

By examining 5mC and 5hmC distribution across gene regions in normal and malignant liver, we found that both 5mC and 5hmC formed a deep dip around TSSs (Supplemental Fig. S7A-B), which is consistent with the previous result of 5hmC depletion in CGI and regions around TSS (Fig. 1A) (Wu and Zhang 2014). However, we further found that in contrast to 5mC, 5hmC exhibited several significant peaks around gene regulatory regions. First, 5hmC formed a bimodal distribution around TSSs of genes. Second, 5hmC level increased along gene body toward 3’ terminus of gene and formed another peak at TTR right after transcriptional termination sites (TTSs) (Supplemental Fig. S7A-B). Although tumors experience global 5hmC loss compared to their normal counterparts, 5hmC enrichment around TSS and TTS still endured in liver tumors, perhaps suggesting a gene regulatory function (Supplemental Fig. S7B).

In line with these results, the distribution of histone modifications around genes strongly associated with the distribution of 5hmC. For example, H3K4me1 associated with 5hmC and was significantly enriched at the flanking regions of TSS (Supplemental Fig. S7C), while H3K4me3 was enriched at TSS where 5hmC depleted (Supplemental Fig. S7D). H3K36me3, which is positively correlated with 5hmC, also showed increasing modification level along gene body toward 3’ end of genes (Supplemental Fig. S7E). Given histone modifications usually coupled with status of gene expression and were not as stable as 5mC (Cedar and Bergman 2009), the strong associations between 5hmC and active histone modifications suggest that 5hmC may be a mark for active genes and can be more dynamically regulated by cells compared to 5mC.

Since CGIs and 5hmC overlap with gene regulatory regions such as the upstream region of TSSs, we further asked whether 5hmC enrichment at gene promoters is dependent on the existence of CGIs and their shores. To address this question we compared 5hmC enrichment at active genes (FPKM > 1) with and without promoter CGI. Compared to genes with promoter CGIs, those genes without promoter CGIs showed almost no 5hmC enrichment around TSS (Supplemental Fig. S7F). This suggests that CGIs and their shores, rather than the location of transcriptional start sites, shape the 5hmC distribution pattern around TSS.

### Promoter, TTR and gene-body hydroxymethylation positively correlated with gene expression

To determine the relationship between 5hmC and gene expression, we generated transcriptome profiles for samples used in this study by RNA-Seq. Based on the 5hmC distribution around genic region, we examined the gene regulatory role of hydroxymethylation at promoter, TTR, and gene body. We classified all genes within tissues into four groups according to their expression levels and found that presence of 5hmC on promoter, gene body, and downstream TTS all show a significant positive correlation with gene expression. The Spearman correlation coefficients (*r*) between 5hmC and gene expression were 0.21, 0.30 and 0.47 for promoter (2 kb upstream of TSS), TTR (2 kb downstream of TTS) and gene body, respectively. Furthermore, compared to inactive genes which have relatively uniform 5hmC distribution throughout genic regions, active genes rather presented two 5hmC peaks in both promoter and TTR, which became more obvious with increased gene expression (Fig. 2A-B).

**Figure 2.**
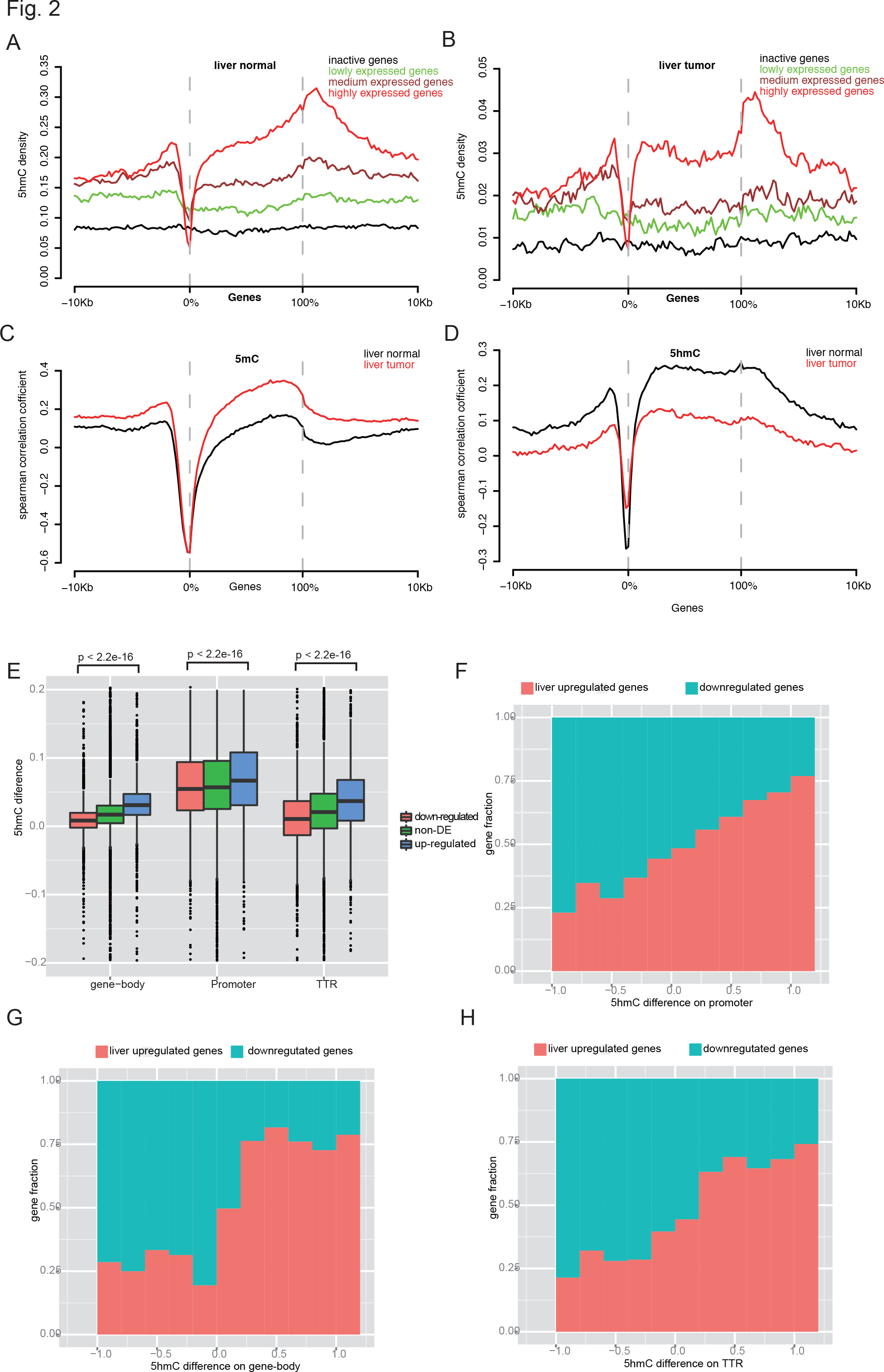
5hmC distribution around genic region and 5hmC positively correlated with gene expression. (A-B) 5hmC distribution across active and inactive genes and their 10 Kb flanking regions in liver normal (A) and liver tumor (B). All genes were classified into active and inactive genes. Active genes were further divided into three groups with equal number of genes by quintile, based on gene expression (FPKM value): lowly, medium, and highly expressed genes. (C-D) Spearman correlation coefficient around genic region between 5mC (C)/5hmC (D) and gene expression. (E) 5hmC difference between normal liver and lung at three genomic features in liver up-regulated, down-regulated and non-DE genes. (F-H) Comparison of 5hmC difference between normal liver and lung on promoter (F), gene body (G) and TTR (H) and gene expression changes between tissues.

We then plotted Spearman correlation coefficients, around genic regions, between 5mC/5hmC and gene expression (Fig. 2C-D), and found a very different pattern between normal and tumor, and between 5mC and 5hmC. For 5mC, except for regions around TSS at which 5mC showed strong negative correlation with gene expression in both normal and tumor tissues, tumors showed positive and much higher correlation coefficients than that in matched normal around genic region. In contrast, for 5hmC, due to extensive loss of 5hmC, tumors showed positive but much lower correlation coefficients across genic regions compared to normal.

We next studied the relationship between level of 5hmC and the difference of gene expression between the normal liver and lung by first identifying differentially expressed (DE) genes. We found 3,771 up-regulated genes and 3,952 down-regulated genes in liver compared to lung, and then calculated the difference in hydroxymethylation level between lung and liver (5hmC_liver_ - 5hmC_lung_) at promoter, TTR, and gene body of those DE genes. Due to much higher global hydroxymethylation level in liver than lung, the average 5hmC level of promoter, TRR, and gene body in liver is higher than that of lung, regardless of whether the genes are up-regulated or down-regulated. However, we found that 5hmC differences at promoter, TRR, and gene body between liver and lung in up-regulated genes are significantly higher than that of down-regulated genes (Wilcoxon test, *p* < 2.2e-16, Fig. 2E). Furthermore, if we divided those DE genes into several groups according to 5hmC difference between tissues, we found that 5hmC difference between liver and lung tissues at promoter, TRR, and gene body showed positive correlation with the fraction of liver up-regulated genes (Fig. 2F-H). Genes in the liver with higher 5hmC level at any of the three gene regulatory region categories were more likely to be up-regulated in liver. In summary, these results reveal a unique 5hmC distribution pattern at genic regions and particular enrichment in active genes and suggest an important role for 5hmC in gene regulation both within tissues and across tissue types.

### 5hmC defines CGI shores and associates with H3K4me1 mark

Our previous studies provided evidence that most t-DMRs and c-DMRs surprisingly overlap with CGI shores but not CGIs, which usually disrupt methylation boundaries of CGI boundaries, and that DNA methylation changes in CGI shores strongly correlate with changes in gene expression (Irizarry et al. 2009). The results in this study also confirmed the previous finding (Supplemental Fig. S8). Here, we examined the 5hmC distribution on CGIs and their shores according to their location and to the expression status of their associated genes. Interestingly, we found that 5hmC is only enriched in CGI shores of active genes, but not with CGI shores of inactive genes or non-promoter CGI shores. 5hmC formed a unique bimodal distribution pattern around CGI shores of active genes in both normal and malignant liver (Fig. 3A-B). We obtained similar results in human stem cells and brain tissue by analyzing a published whole genome TAB-Seq 5hmC datasets (Supplemental Fig. S9A-B), supporting a universal 5hmC bimodal distribution pattern around CGI shores of expressed genes regardless of tissue type and disease status.

**Figure 3.**
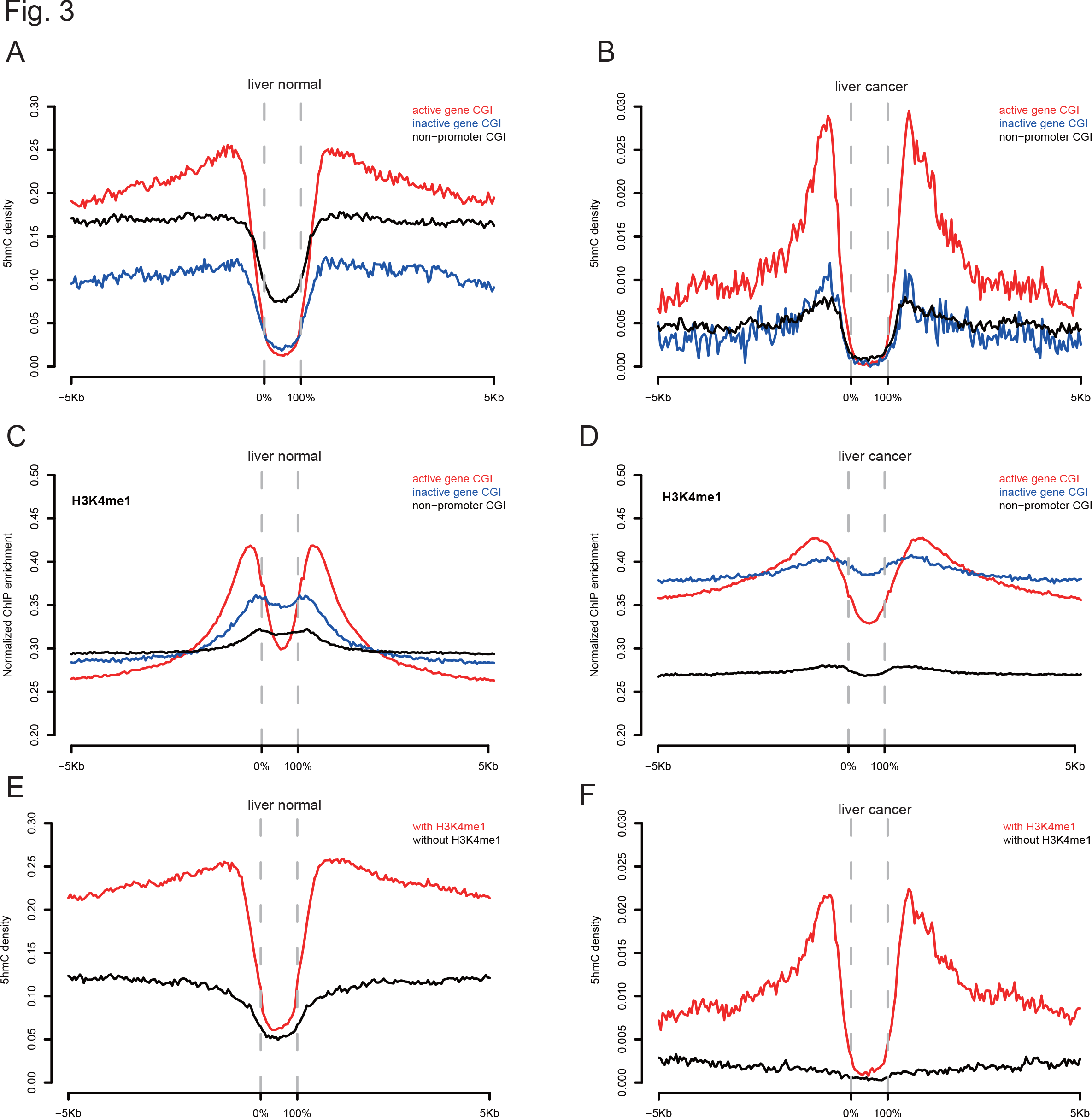
5hmC showed bimodal distribution around CGIs and associated with H3K4me1. (A-B) 5hmC distribution around different CGI categories in live normal (A) and liver tumor (B). (C-D) H3K4me1 enrichment around different CGI categories in live normal (C) and liver tumor (D). (E-F) H3K4me1 associated with 5hmC enrichment in CGI shores in liver normal (E) and liver tumor (F).

To gain insight into the potential functional role of 5hmC in CGI shores, we analyzed the association of 5hmC distribution with a variety of active histone modification marks using publicly available ChIP-Seq data (Roadmap Epigenomics et al. 2015). While H3K4me3, H3K9ac and H3K27ac showed peaks at CGI of active genes (Supplemental Fig. S9C-H), H3K4me1 showed bimodal peaks around CGI shores similar to that of the 5hmC (Fig. 3C-D), suggesting potential crosstalk between H3K4me1 and 5hmC at CGI shores. To thoroughly examine the association between 5hmC and H3K4me1, we divided all CGI shores into two groups—with or without H3K4me1 modification—and found that only CGI shores with H3K4me1 modification showed clear 5hmC bimodal peaks (Fig. 3E-F). CGIs with H3K4me1 on shores also showed H3K4me3 modification that tightly associated with active genes, again indicating positive correlation between 5hmC/H3K4me1 and gene expression (Supplemental Fig. 10). Taken together, these results suggest that 5hmC and the association with H3K4me1 mark contribute to the function of CGI shores in regulating gene expression.

### 5hmC correlates positively with H3K4m1 and negatively with 5mC in both t-DMRs and c-DMRs

Consistent with the observation that liver showed a lower global 5mC methylation level than lung (62.08% vs. 65.32%, *t* test, *p*-value=0.01354), we identified more hypomethylated than hypermethylated t-DMRs in normal liver (Supplemental Table S7-8) for 5mC. We divided all t-DMRs into three groups: CGI-associated, CGI shore-associated, and other DMRs. Consistent with our previous study (Irizarry et al. 2009), very few t-DMRs were CGI-associated (2.9%) while most t-DMRs were CGI shore-associated (21.0%) or distant from CGI (76.1%), the latter mostly within gene bodies (66.5%).

Our previous study showed that c-DMRs highly enriched for overlap with t-DMRs (Irizarry et al. 2009). Following this point, we further found that liver hypermethylated c-DMRs significantly overlapped with t-DMRs where liver was comparatively hypermethylated than that of liver hypomethylated (13.4% vs. 2.1%, chi-square test, *p* < 2.2e-16, Supplemental Table S9). Similarly, lung hypermethylated c-DMRs significantly overlapped with t-DMRs where lung was comparatively hypermethylated than that of lung hypomethylated (14.8% vs. 1.8%, chi-square test, *p* < 2.2e-16, Supplemental Table S9). In line with this result, liver cancer down-regulated genes significantly enriched in liver up-regulated genes between normal tissues than that of liver down-regulated genes (42% vs. 6.1%, chi-square test, *p* < 2.2e-16). Similar result was also observed in lung cancer (54.2% vs. 8.8%, chi-square test, *p* < 2.2e-16). Taken together, these results suggested that the aberrant DNA hypermethylation might disrupt normal tissue-specific methylation landscape during tumorigenesis and result in transcriptional silencing of genes that usually actively expressed in normal tissues.

To determine the association between t-DMR and 5hmC, we overlapped t-DMRs with identified liver-and lung-specific 5hmC regions (Fig. 4A). We found that in general liver-hypermethylated t-DMRs were hypo-hydroxymethylated, while liver-hypomethylated t-DMRs were hyper-hydroxymethylated between normal liver and lung (5hmC_liver_ - 5hmC_lung_, −0.014 vs. 0.048, Wilcoxon test, *p*-value < 2.2e-16), indicating a negative correlation between 5mC difference and 5hmC difference on t-DMRs (Fig. 4B). Furthermore, we found that DNA methylation changes at t-DMRs also negatively correlated with H3K4me1 difference (Fig. 4C). Consistently, t-DMRs where liver was comparatively hypermethylated overlapped with lung-specific H3K4me1 peaks, while t-DMRs where liver was comparatively hypomethylated overlapped with liver-specific H3K4me1 peaks (Supplemental Table S7-8). These results suggested strong associations among dynamic changes of 5mC, 5hmC and H3K4me1, and more importantly, these H3K4me1-associated 5mC and 5hmC changes corresponded to changes in gene expression between tissue types as shown above.

**Figure 4.**
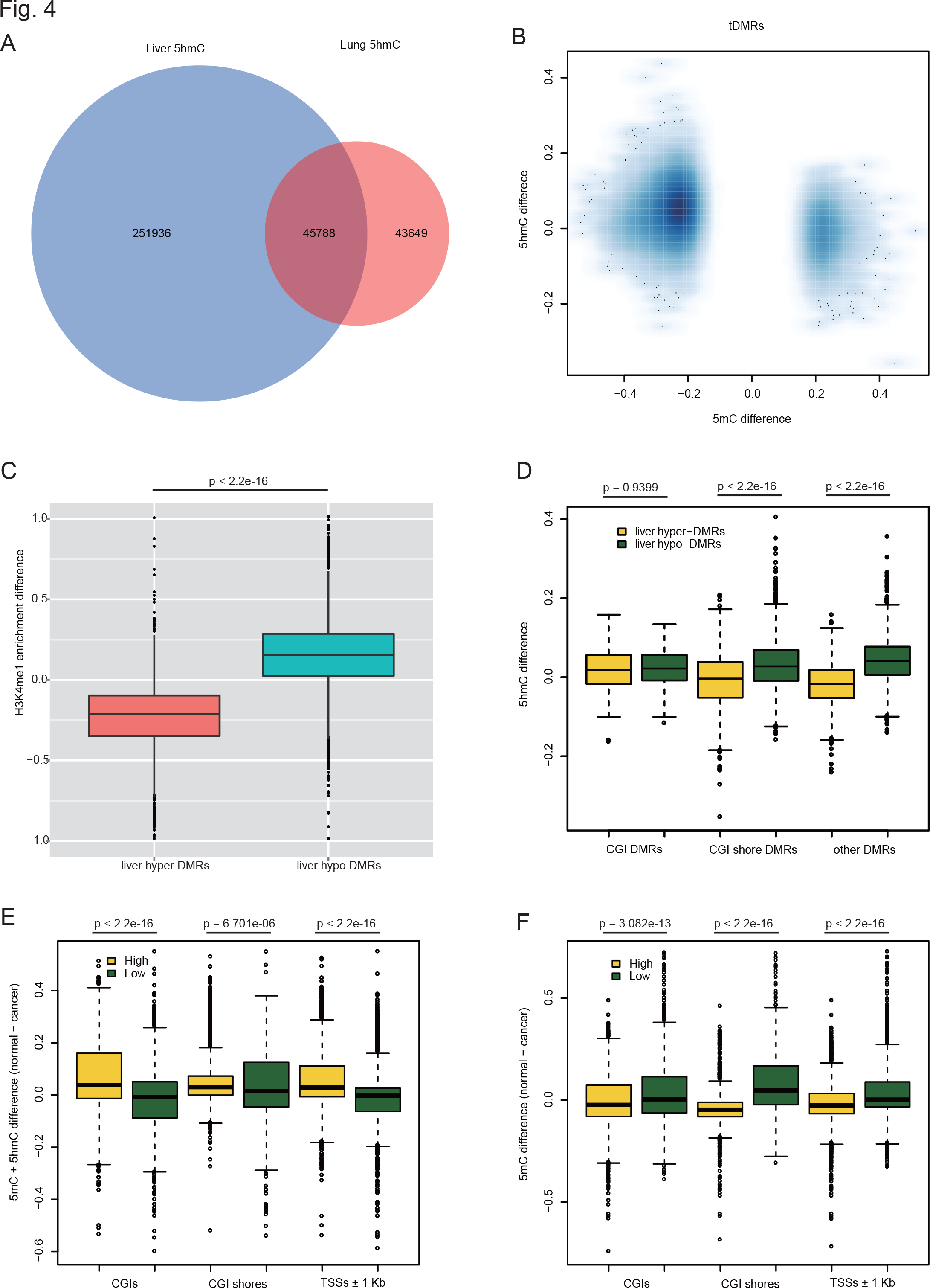
5mC change of t-DMRs and c-DMRs anti-correlated with 5hmC and H3K4me1 changes. (A) Venn diagram of identified liver and lung 5hmC regions. (B) Negative correlation between 5mC different and 5hmC difference on t-DMRs. (C) Negative correlation between 5mC different and H4K4me1 enrichment difference on t-DMRs. (D) 5hmC difference on different t-DMR categories. 5hmC difference on non-CGI t-DMRs showed larger difference than that of CGI t-DMRs. (E) Cytosine modification difference (5mC + 5hmC) between normal and tumor at promoters with high and low 5hmC level in liver (F) Only DNA methylation difference (5mC) between normal and tumor at promoters with high and low 5hmC level in liver. Promoters were ranked according to 5hmC level and top and lowest 10th percentiles of promoter were used for our analysis.

Next we explored c-DMRs in two types of cancer (Supplemental Table S9-11). We examined the association between changes in 5hmC and 5mC at c-DMRs. Similar to t-DMRs, 5hmC change in c-DMRs also showed strong negative correlation with 5mC change (Fig. 5A-B and Supplemental Fig. S11A-B). In other words, loss of 5hmC corresponded to gain of 5mC and vice versa. For both t-DMRs and c-DMRs, we found that the negative correlation between 5hmC and 5mC changes is stronger in CGI shore-associated DMRs and other-DMR categories than CGI-associated DMRs (Fig. 4D] and Supplemental Fig. S11C-D), which is consistent with depletion of 5hmC in CGIs. Again, similar to t-DMRs, the negative correlation between 5mC and H3K4me1 was also observed in c-DMRs (Supplemental Fig. S11E-F). Taken together, our results suggested that gene expression pattern in different tissues could be regulated and achieved through intricate interplays among DNA methylation, hydroxymethylation, and histone modification such as H3K4me1.

**Figure 5.**
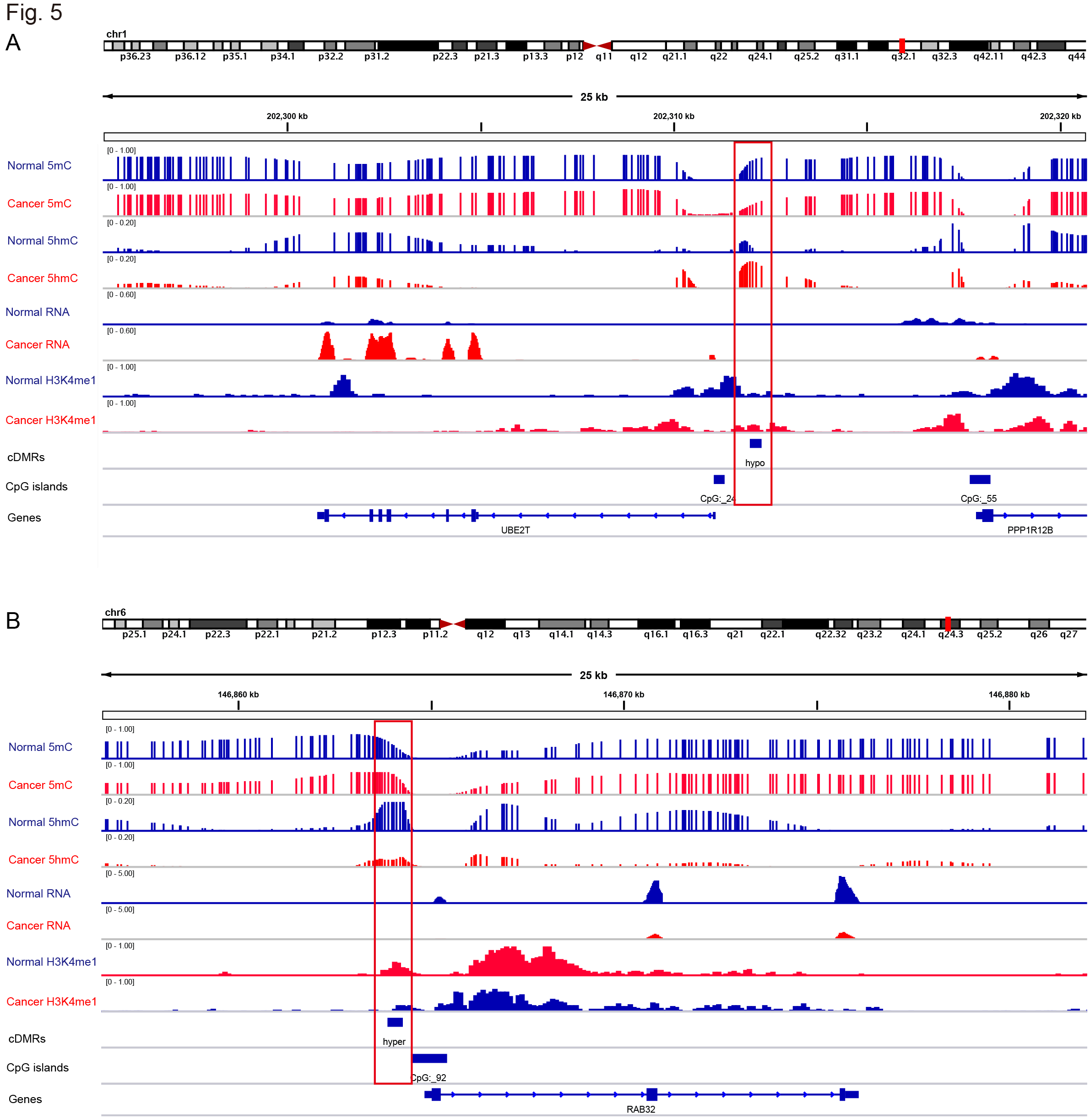
Two examples of liver c-DMRs that showed 5mC changes was regulated through 5hmC and associated with gene expression changes. 5mC, 5hmC and gene expression level of both liver normal (blue tracks) and cancer (red tracks) were displayed. Red boxes indicate the location of c-DMRs. (A) A hypomethylated liver c-DMR located in CpG island shore near the promoter of UBE2T gene. (B) A hypermethylated c-DMR located in CpG island shore near the promoter of RAB32 gene.

The negative correlation between 5mC and 5hmC observed here is not consistent with the finding, in a previous study using colon tissue, that high 5hmC promoters in normal tissue are prone to loss of DNA methylation in tumors thus resistant to DNA hypermethylation in cancer (Uribe-Lewis et al. 2015). However, that study used Infinium27K array to obtain methylation profiles, which cannot distinguish 5mC and 5hmC. We performed similar analysis using our data and found that promoters with high 5hmC level in normal indeed tend to lose DNA methylation in cancer, compared to that of low 5hmC promoters when considering 5mC and 5hmC together as DNA methylation differences (Fig. 4E and Supplemental Fig. S11G). However, when removing the effect of 5hmC and only using 5mC, we found the opposite pattern, that promoters with high 5hmC level in normal in fact tend to be hypermethylated in cancer, whereas low 5hmC promoters tend to be hypomethylated (Fig. 4F and Supplemental Fig. S11H). This result suggests that high 5hmC level maintain promoter hypomethylation in normal, and that global loss of 5hmC could result in hypermethylation at promoter and dysregulation of gene expression in cancer. This result is consistent with the observation that there are more small hypermethylated c-DMRs than hypomethylated c-DMRs, and again confirmed the anti-correlation between 5hmC and 5mC. This result also demonstrated the importance of distinguishing 5mC and 5hmC in DNA methylation analysis, especially for those tissues with relative high 5hmC level, and the conclusions from previous studies using traditional methods may need to be reassessed. That said, it is quite possible that the colon differs from other tissues, particularly given its low level of 5hmC in normal, although one would need to use a method distinguishing 5hmC from 5mC in the 5mC assays, as done here.

## Discussion

In this study, we have performed base-level resolution analysis of 5mC and 5hmC in normal and matching malignant tissues from human liver and lung. Our results revealed that 5hmC is an important epigenetic mark of active genes that is strongly associated with active histone modifications, and could play a role at gene expression mediated by DNA demethylation.

Interestingly, our results showed 5hmC is significantly enriched in CGI shore relative to CGI itself, and revealed an interesting 5hmC bimodal distribution pattern around CGI.

Supporting this finding, recent studies using hmeDIP-Seq showed that 5hmC is enriched within the shores of promoter CpG islands in human normal colon tissues and embryonic carcinoma cell line (Putiri et al. 2014; Uribe-Lewis et al. 2015). By overlaying these data with RNA-Seq of the same tissues, we found that 5hmC is only enriched in CGI shores which are associated with active genes, and not with CGI shores which are associated with inactive genes or with nonpromoter associated CGI shores. In addition, this unique 5hmC distribution around CGI also coupled with H3K4me1. Although our previous studies have reported that DNA methylation in CGI shore significantly regulates gene expression across tissues (Irizarry et al. 2009; Hansen et al. 2011), the underlying molecular mechanism remains unclear. Our results demonstrate that CGIs and their shores function as important regulatory regions that are enriched with variety of histone modification, and these histone modifications around CGIs and their shores could help them be subjected to regulation of TET-mediated DNA methylation.

Besides 5hmC bimodal peaks around TSSs, 5hmC level increased along gene bodies and formed another peak right after transcriptional termination sites (TTSs). Strikingly, the 5hmC level is even higher at TTSs than at promoter regions and also positively correlated with gene expression, suggesting the possibility that 5hmC around TTSs could affect gene expression by interacting with transcription termination factors and regulating RNA polymerase II (Pol II) processivity. Supporting this idea, an *in vitro* study showed DNA templates incorporated with 5fC and 5caC, oxidation variants of 5hmC, can dramatically reduce the transcription rate of Pol II and stall Pol II (Kellinger et al. 2012). Whether 5hmc enrichment at TTS contributes to oxidation variants that give rise to transcription termination *in vivo* requires further detailed investigation.

Our results further showed that many t-DMRs and c-DMRs were located in gene body and also significantly affected gene expression between tissues. Furthermore, those DMRs usually overlap with tissue-specific 5hmC and H3K4me1 peaks, which usually associate with distal regulatory elements and is used for defining enhancers. In additions, previous studies showed that intragenic enhancers within gene body can regulate gene expression by acting as alternative promoters (Kowalczyk et al. 2012). Recent work also showed that extensive loss of 5hmC at enhancers mediated by Tet2 deletion was coupled with enhancer hypermethylation and affected gene expression during early stage of mouse stem cell differentiation (Hon et al. 2014). Taken together, these results suggest that dynamic change in 5mC/5hmC at intragenic enhancer regions could be an important epigenetic mechanism in regulation of gene expression during tissue differentiation and tumorigenesis.

In cancer compared to normal, 5hmC was quantitatively diminished (~70%) and partially retained at regulatory regions. Moreover the difference in 5hmC between euchromatic and heterochromatic regions in normal tissue is essentially lost in cancer. Similarly, the specific relationship between 5hmC and chromatin marks in normal tissue is largely erased in tumors. These results suggested that 5hmC landscape change in cancer could associate with chromatin structure change and regulation of gene expression during tumorigenesis.

These experiments also showed a strong anti-correlation between 5hmC and 5mC changes in both t-DMRs and c-DMRs, with this effect stronger in CGI shores. Global loss of 5hmC is considered to be a hallmark of cancer cells, and impairment in the TET-mediated DNA demethylation machinery is described in several tumor types (Delhommeau et al. 2009; Langemeijer et al. 2009; Dang et al. 2010; Shibata et al. 2011). Considering that TET enzymes preferentially bind CpG enriched region, such as CGIs, dysfunction of TETs-mediated DNA demethylation in cancer could explain why more hypermethylated c-DMRs are located in CGIs, which is also supported by previous result that loss of 5hmC in TET-depleted human ECC line coincides with genes susceptible to aberrant hypermethylation.

The antagonistic role of 5mC and 5hmC in normal tissue, and the bimodal distribution of 5hmC at TSSs and at H3K4Me1 sites associated with enhancers, suggests that DNA methylation is mediated by the balance and topological organization of 5mC and 5hmC, creating a mechanism for both stable gene expression but also substantial and abrupt changes during normal development. Finally, the DNA methylation maintenance machinery is robust and self-sustained. In agreement with this, *DNMT1* mutations are rarely observed in cancer. Therefore, we postulate that disruption of 5hmC could lead to instability of methylation marks that are then selected for and maintained by the *DNMT1* machinery to increase the proliferative advantage of cancer cells over normal.

Based on our result, the majority of DNA loss in tumors seems to be due to passive demethylation, especially in large hypomethylated blocks where 5hmC is depleted. However, we also found both hyper-and hypo-methylated c-DMRs at CGI shores where 5hmC is significantly enrich and associates with bimodal H3K4me1. Therefore, it is possible that active demethylation plays an important role in those particular regulatory elements. Future studies would be necessary in order to determine whether DNA demethylation is passive or active in tumorigenesis. For example, dynamic 5hmc profiling during different stages of tumorigenesis or during induced tumorigenesis in the absence of TET enzymes.

## Methods

### Preparation of hydroxymethylated lambda phage genome

1 μg unmethylated lambda DNA (Promega) was treated with SssI methyltransferase (Zymo) overnight and cleaned up using Genomic DNA Clean & Concentrator (Zymo). To make sure all CpG sites of lambda genome were fully methylated, SssI treatment and clean-up were repeated. Then 500 ng CpG methylated lambda DNA was bisulfite converted using EZ DNA Methylation-Lightning Kit following manufacturer’s manual (Zymo). 10 ng bisulfite converted lambda DNA from above step was used to perform whole genome amplification by GenoMatrix Whole Genome Amplification Kit (Active Motif) using 5-hydroxymethylcytosine dNTP Mix (Zymo) instead of dNTP mix in the kit. To achieve as high hydroxymethylation level as possible, PCR reaction of whole genome amplification was repeated for another 3 times using 5-hydroxymethylcytosine dNTP and PCR product from last time as template every time.

### oxBS-Seq and BS-Seq library preparation

Genomic DNA was extracted using DNeasy Blood & Tissue Kit (Qiagen). 4 μg genomic DNA plus spike-in 20 ng hydroxymethylated lambda phage and non-methylated *E. coli* DNA control (Zymo) in 150 μl water was sheared into ~10 kb fragments using g-Tube (Covaris) following manufacturer’s instructions. And sheared DNA was cleaned up using GeneJET PCR Purification Kit (Thermo Scientific) with a modified protocol (Protocol 03, TrueMethyl Preparation of High Molecular Weight gDNA, http://www.cambridge-epigenetix.com/en_US/resources/application-notes). Then oxidative bisulfite (oxBS) and only bisulfite (BS) converted DNA templates were generated using TrueMethyl 24 Kit (Cambridge Epigenetix) for each sample according to manufacturer’s instructions. Last, oxBS and BS converted DNA were used to construct oxBS-Seq and corresponding BS-Seq libraries using EpiGenome Methyl-Seq Kit (Epicentre) following manufacturer’s instructions.

### oxBS-Seq and BS-Seq data processing

Paired-end HiSeq2000 sequencing reads from oxBS-Seq and BS-Seq were aligned by BSmooth bisulfite alignment pipeline (version 0.7.1) (Hansen et al. 2012) as previously described in detail (Hansen et al. 2011). Briefly, reads were aligned by Bowtie2 (version 2.0.1) against human genome (hg19) as well as the lambda phage and *E. coli* genomes. After alignment, methylation measurements for each CpG were extracted from aligned reads. We filtered out measurements with mapping quality < 20 or nucleotide base quality on cytosine position < 10 and we also removed measurements from the 5’ most 7 nucleotides of both mates. Methylation level of lambda phage and *E. coli* genomes were used to access oxidation and bisulfite conversion rates respectively.

### t-DMR and c-DMR identification

To identify t-DMRs and c-DMRs that were only contributed by 5mC and do not involve 5hmC changes, oxBS-Seq data were used for DMRs identification by bsseq package, which can borrow statistical power from neighboring CpG sites and biological replicates and was successfully applied to DMRs identification even with low sequencing coverage in our previous studies (Hansen et al. 2011; Hansen et al. 2014). We reasoned that oxBS data, measuring only 5mC should behave similarly to standard WGBS data measuring the sum of 5mC and 5hmC and we therefore applied BSmooth with standard parameters. Specifically, for small DMRs we used a smooth window containing either 70 CpGs or a width of 1kb, whichever is larger and for large DMRs (blocks) we used a smoothing window containing either 500 CpGs or a width of 20kb, whichever is larger. Following smoothing, putative DMRs were identified using a t-stat cutoff (see below). For this computation, we only considered CpGs with coverage of at least 5 in at least 2 samples in each group, and we only considered putative DMRs with a methylation difference of at least 20% (small DMRs) or 10% (large DMRs) and a length greater than 5kb. These arbitrarily chosen cutoffs are more stringent that what we have employed in earlier work (Hansen et al. 2011; Hansen et al. 2012; Hansen et al. 2014). Final DMRs depends on both the t-statistic cutoff and the procedure for assigning significance. Previously we employed a very stringent permutation procedure, which controls the family-wise error rate; this is a much more stringent error rate than the widely used false discovery rate (FDR) (Hansen et al. 2014). In this work, we approached the problem differently. We systematically tested a range of t-statistic cutoffs (1.65, 2, 2.5, 3, 3.5, 4, 4.6) and for each cutoff we found putative DMRs by as consecutive sets of CpGs with a t-statistics exceeding the cutoff. We compute p-values associated with each DMR using the following procedure. First we compute a CpG level p-value based on a standard t-test for the smoothed methylation data using an asymptotic reference distribution. Next, these CpG level p-values were combined into a region level p-value by using a modified Stouffer-Liptak method as implemented in the Comb-p software (Pedersen et al. 2012). Next, these regional level p-values were corrected for multiple testing and we kept regions with an FDR less than 5%. This gives us a set of DMRs for each t-statistic cutoff. We then choose the cutoff that yielded the most differentially methylated regions, effectively optimizing empirical power (Supplementary Figure 12). To assess the error rate we permuted the sample labels and repeated the procedure (Supplementary Figure 12) yielding almost no DMRs with an FDR less than 5%, showing our low error rate.

### 5hmC region and site identification

According to the principle of oxidation bisulfite sequencing, hydroxymethylation can be ascertained by the methylation difference of oxBS-Seq and corresponding BS-Seq data for each sample. Since hydroxymethylation level is usually very low in human tissues except for brain tissues and it needs high sequencing depth to get accurate measurement, we used a similar smooth-based algorithm like above DMRs identification in this study to identify hydroxymethylation regions (5hmC regions). Effectively, this is identifying differences between BS-Seq and oxBS-Seq libraries. We smoothed both data types using default parameters from BSmooth (as above, a window size encompassing at least 1kb or 70 CpGs). Assessment of significance was done as described above (Supplemental Fig. 12C-D). After obtaining 5hmC regions, only CpGs passing the *t*-statistic threshold within 5hmC regions were considered as hydroxymethycytosines (5hmC sites) and used for analysis in this study. In this study, 5hmC density is estimated by the proportion of 5hmC sites out of all CpGs within a certain genomic feature/region. 5hmC level for CpG sites or regions is estimated by the subtraction between BS-Seq and oxBS-Seq libraries based on smoothed values.

### c-DMRs and 5hmC regions replications

Another liver cohort including 6 normal-cancer pairs was kindly provided by Professor Robert Albert Anders. Genomic DNA was purified using DNeasy Blood & Tissue Kit (Qiagen). 1.5 μg DNA for each sample was used to generate oxidative bisulfite (oxBS) and only bisulfite (BS) converted DNA templates by TrueMethyl 24 Kit (Cambridge Epigenetix) following manufacturer’s instructions. Primers for bisulfite sequencing PCR were designed using MethPrimer (http://www.urogene.org/methprimer/) and sequences of all primers were listed in Supplemental Table S12. Locus-specific PCRs were performed by nested PCR using both oxBS-and BS-converted DNA as templates. Then all amplicons from same sample were pooled, and barcoded libraries were prepared with TruSeq DNA PCR-Free Library Preparation Kit (Illumina) following manufacturer’s instruction. Amplicon sequencing was performed on a MiSeq instrument (Illumina).

### RNA-Seq library preparation and data processing

Total RNA was extracted using RNeasy Mini Kit (Qiagen). RNA-Seq libraries were constructed using TruSeq Stranded mRNA LT Sample Prep Kit (Illumina) according to manufacturer’s manual.

Paired-end HiSeq2000 sequencing reads were aligned against human genome (hg19) by OSA (version 2.0.1) (Hu et al. 2012) with default parameters. After alignment, only uniquely aligned reads were kept for further analysis. Gene annotation information was downloaded from GENCODE (http://www.gencodegenes.org/releases/19.html, release 19). Reads count for each gene of all samples were estimated using HTSeq (http://www-huber.embl.de/users/anders/HTSeq/doc/overview.html) and then were used to identify differentially expressed (DE) genes using DESeq2 package (Love et al. 2014). Genes with FDR < 0.01 and fold-change > 2 between groups were considered as DE genes.

### ChIP-Seq data processing

Uniformly processed ChIP-Seq data used in this study including normal and tumor tissues of human liver and lung were download from Roadmap Epigenomics Project (http://www.broadinstitute.org/~anshul/projects/roadmap/mapped/stdnames30M/).

For narrow histone modification peaks including H3K4Me1, H3K4Me3, H3K9Ac, and H3K27ac, MACS2 were used for peak calling with default parameters (Zhang et al. 2008). While for broad histone modifications peaks in including H3K27Me3, H3K36me3, H3K9Me3, large domains were identified using RSEG which is based on hidden Markov model (HMM) and specifically designed for identifying broad histone peaks (Song and Smith 2011).

Based on ChIP-Seq, enhancers were defined as H3K4me1 peaks that are at least 2 kb away from any transcriptional stat site of annotated genes. Enhancers overlapped with H3K27ac peaks were defined as active enhancers, while others were poised enhancers.

To plot each histone modification on CpG islands (CGIs) and their flanking regions, we divided flanking sequences into bins with fixed length (in bp) and CGIs themselves into bins with fixed percentage of each length. ChIP enrichment was measured and normalized using a previous published method (Hawkins et al. 2010). In brief, the number of reads per kilobase of bin per million reads sequenced (RPKM) was calculated for each ChIP and its input control (denoted as RPKM_chIP_ and RPKM_input_). ChIP enrichment is measured as ΔRPKM = RPKM_ChIP_ − RPKM_input_ and ChIP enrichment regions should have ΔRPKM > 0. Then all ΔRPKM were normalized to a scale between 0 and 1 and the average normalized ChIP enrichment signals across all bins were plotted for each histone marks.

## Data access

Sequence data generated in this study has been submitted to NCBI Gene Expression Omnibus (GEO) under accession numbers GSE70090 (oxBS-Seq and BS-Seq) and GSE70089 (RNA-Seq).

## Acknowledgements

We thank Rebecca Curley, Rakel Tryggvadottir, Birna Berndsen and Adrian Idrizi for guidance in library construction and sequencing assistance. This work was supported by NIH Grant CA54358 to Andrew Feinberg.

## Author contributions

X.L. and A.F. designed the research and wrote the manuscript; X.L., Y.L. and T.S. performed the experiments. X.L. performed the analysis, K.D.H provided substantial statistical oversight and writing regarding our improved BSmooth method, and Y.L. reviewed the accuracy of the statistical code and analysis.

## Supplemental Figure Legends

**Supplemental Figure 1.** Global methylation and hydroxymethylation levels in samples used in this study. (A) methylation (B) hydroxymethylation. Smoothed methylation and hydroxymethylation levels of all CpG sites across genome was averaged as global 5mC and 5hmC levels.

**Supplemental Figure 2.** Verification of BSmooth algorithm. High whole-genome Spearman correlation between low and high coverage datasets for (A) 5mC (B) 5hmC. 5hmC level of CpG sites were estimated based on smoothed value. (C) Close agreement of methylation profiles between low and high coverage data. Each grey dot represents methylation level of a CpG site estimated by 55 × dataset. Black (15 ×) and red lines (55 ×) represent methylation levels estimated based on smoothed profiles. (D) 5mC profiles of both oxBS-Seq and BS-Seq libraries and 5hmC profiles between low and high coverage data on an identified 5hmC region. Identified 5hmC regions were indicated as grey boxes, while CGIs were indicated as black boxes.

**Supplemental Figure 3.** Liver c-DMRs replications using another cohort. Boxplot were plotted for amplified c-DMR. The blue dots represent average methylation levels of CpGs within c-DMRs for each normal/cancer group. Paired *t* test was performed for each c-DMRs between normal and cancer groups and *p*-value was showed in upper-right of each panel.

**Supplemental Figure 4.** Liver normal 5hmC replications using another cohort. 5mC levels of BS-Seq (left) and oxBS-Seq (right) libraries were plotted for each 5hmC region. Paired *t* test was performed for each 5hmC region between BS-Seq and oxBS-Seq groups and *p*-value was showed in upper-right of each panel.

**Supplemental Figure 5.** Genome-wide correlation between 5hmC and histone modifications. (A) liver normal, (B) liver tumor (C) lung normal, (D) lung tumor. Whole genomes were divided into 5 Kb sliding windows. 5hmC level and histone modification enrichments were calculated for each window and used for estimating Spearman correlation coefficients among them.

**Supplemental Figure 6.** 5hmC fold enrichment on c-DMRs and t-DMRs. (A) 5hmC fold enrichment of liver c-DMRs (B) and lung c-DMRs. (C) 5hmC fold enrichment of t-DMRs in liver and lung. Y-axis were log scaled. 5hmC enrichment fold in each categories was calculated as the ratio of 5hmC density of that feature to genome-wide average 5hmC density.

**Supplemental Figure 7.** Distribution of 5mC, 5hmC, and histone distribution across genes and their 10 Kb flanking regions. 5mC and 5hmC distribution around genic region (A) liver normal (B) liver tumor. (C) H3K4me1 and H3K9ac in liver normal. (D) H3K27ac and H3K4me3 in liver normal. (E) H3K36me3 in liver normal (F) 5hmC distribution around TSS for active genes with promoter CGI or without in liver normal.

**Supplemental Figure 8.** Gene expression changes is correlated with both t-DMRs and c-DMRs at CGIs and CGI shores. (A) t-DMRs between normal liver and lung (B) liver c-DMRs (C) lung c-DMRs. Y-axis indicates log2 ratio of gene expression (measured by FPKM) between tissues. DMRs that are located within 2 kb from TSSs of genes were used for plotting. Block circles represent CGI shore-associated DMRs, while red circles represent CGI-associated DMRs.

**Supplemental Figure 9.** 5hmC distribution and histone modification enrichment around different CGI categories. (A-B) 5hmC distribution around different CGI groups in human stem cell (A) and fetal brain (B). (C-H) Histone modification enrichment around different CGI groups in liver normal (C, E and G) and liver tumor (D, F and H).

**Supplemental Figure 10.** H3K4me3 enrichment around CGIs with and without H3K1me1 peaks. (A) liver normal (B) liver cancer.

**Supplemental Figure 11.** 5mC change anti-correlated with 5hmC change on c-DMRs. (A-B) Scatter plots between 5mC difference and 5hmC difference in liver c-DMRs (A) and lung c-DMRs (B). (C-D) Boxplots of 5hmC difference in different liver c-DMRs (C) and lung c-DMRs (D) categories. (E-F) Boxplots of H3K4me1 modification difference in different liver c-DMRs (E) and lung c-DMRs (F) categories. (G) Cytosine modification difference (5mC + 5hmC) between lung normal and lung tumor at promoters with high and low 5hmC level. (H) Only DNA methylation difference (5mC) between lung normal and lung tumor at promoters with high and low 5hmC level. Promoters were ranked according to 5hmC level and top and lowest 10th percentiles of promoter were used for our analysis.

**Supplemental Figure 12.** *t*-statistic cutoffs were determined to achieve balance of sensitivity and specificity. The numbers of CpGs within identified c-DMRs/5hmCs varies among different chosen cutoffs. The *t*-statistic cutoff selected was the one achieving the largest number of CpGs within the identified DMRs/5hmCs. Sample labels were permuted 2 times and almost no DMRs/5hmC regions were observed in the permutations no matter what cutoff was chosen. (A) liver c-DMRs identification (B) lung c-DMRs identification (C) liver 5hmC region identification (D) lung 5hmC region identification.

